# Biological activity of *Ajuga iva* extracts against the African cotton leafworm *Spodoptera littoralis*

**DOI:** 10.1101/2020.03.12.988428

**Authors:** Leena Taha-Salaime, Galina Lebedev, Jackline Abo-Nassar, Sally Marzouk, Moshe Inbar, Murad Ghanim, Radi Aly

## Abstract

The African cotton leafworm *Spodoptera littoralis*, a major crop pest worldwide, is controlled by chemical insecticides, leading to serious resistance problems. *Ajuga* plants contain phytoecdysteroids (analogs of arthropod steroid hormones that regulate metamorphoses) and clerodanes (diterpenoids exhibiting antifeedant activity). We analyzed phytoecdysteroids and clerodanes in leaf extracts of the Israeli *Ajuga iva* by LC-TOF-MS and TLC, and their efficiency at reducing *S. littoralis* fitness. Castor bean leaves were smeared with an aqueous suspension of dried methanolic crude extract of phytoecdysteroid and clerodanes from *A. iva* leaves (50, 100 and 250 µg/µl). First and third instars of *S. littoralis* larvae were fed with 1 treated leaf for 3 and 4 days, respectively. Mortality, larval weight gain, relative growth rate and survival were compared to feeding on control leaves. To evaluate and localize *A. iva* crude leaf extract activity in the insect gut, we used DAPI and phalloidin staining. Crude extract of *A. iva* leaves (50, 100 and 250 µg/µl) significantly increased mortality of first instar *S. littoralis* larvae (36%, 70% and 87%, respectively) compared to controls (6%). Third instar larval weight gain decreased significantly (by 52%, 44% and 30%, respectively), as did relative growth rate (–0.05 g/g day, compared to the relevant controls). *S. littoralis* larvae were further affected at later stages, with few survivors. Insect-gut staining showed that 250 µg/µl crude leaf extract reduces gut size, with relocation of nuclei and abnormal actin-filament organization. Our results demonstrate the potential of *A. iva* extract for alternative, environmentally safe insect-pest control.

**Key Message:** Insects cause severe damage to numerous crops and their control relies on pesticides. Green control is becoming increasingly popular due to concerns about the negative impacts of pesticides on the environment. Phytoecdysteroids are found in Ajuga plants and affect a wide range of insects at very low concentrations. Here we demonstrate that crude extract from *Ajuga iva* alters the development of *Spodoptera littoralis*. Phytoecdysteroids may therefore be beneficial in IPM programs.

## Introduction

The African cotton leafworm *Spodoptera littoralis* is considered one of the most serious pests of cotton, maize, rice, alfalfa, potato, tomato, ornamentals and orchard trees (Martinez and van Emden 2001). It feeds year-round on the leaves of numerous old- and new-world plant species (Adel El-Sayed et al. 2011). Today, insect pests are mainly controlled by insecticides, which constitute a risk to human health and the environment (Horowitz et al. 2005). Many organic insecticides have been derived from plant sources, and some, such as alkaloids, terpenoids, phenols and steroids, exhibit very high toxicity against a variety of agricultural pests. In this study, we examined the potential use of phytoecdysteroids and clerodanes extracted from *Ajuga* (Lamiaceae) plants to control the African cotton leafworm.

Phytoecdysteroids are plant-produced steroids that are analogs of the steroid hormones that control molting and metamorphosis in arthropods (Dinan 2001). Phytoecdysteroids are present in 5–6% of plant species (Sandlund et al. 2018), generally at higher concentrations than those typically found in arthropods (Dinan 1995a). Most of them possess a cholest-7-en-6-one carbon skeleton (C27), and are synthesized from phytosterols in the cytosol through the mevalonic acid pathway (Dinan 2001). They can mimic insect 20-hydroxyecdysteroid, bind insect ecdysone receptors and elicit the same responses (Sadek 2003). Phytoecdysteroids may cause abnormal larval development, feeding deterrence and ultimately, death (Sadek 2003). Ecdysteroids are not toxic to mammals because their structure is quite different from mammalian steroids, and they are not expected to bind to vertebrate steroid receptors (Lafont and Dinan 2003).

Ecdysone, a natural molting hormone of insects derived from enzymatic modification of cholesterol by p450 enzymes (Dinan 1989), controls developmental events by changing the levels of other ecdysteroids (Lafont 1997). The ecdysone receptor is a nuclear receptor (a ligand-activated transcription factor) that binds to and is activated by ecdysteroids. In *Manduca sexta* larvae, 20-hydroxyecdysone is primarily produced in the prothoracic gland, gut and fat bodies (Grieneisen et al. 1991) from dietary cholesterol, and acts through the ecdysone receptor (Thummel and Chory 2002). In addition, the ecdysone receptor controls development, and contributes to other processes (such as reproduction) (Riddiford et al. 2000), and to interactions between the cytoskeleton (the effector of cell movement and changes in cell shape) and changes in the distribution of actin staining and microfilaments (Otey et al. 1990).

Discovery of the same molecules (phytoecdysteroids) in several plant species suggests that they may be effective against insect herbivores by acting as antifeedants and/or disrupting the insects’ endogenous endocrine levels (Blackford et al. 1996; Dinan 2001; Belles and Piulachs 2014). Low concentrations (2–25 ppm) of phytoecdysteroids deter some insects, while others are resistant to even very high concentrations (400–1000 ppm) (Blackford et al. 1996). Kubo (1997) reported that an extract of *Ajuga remota* containing 20-hydroxyecdysone and cyasterone, added to the diet of *Bombyx mori*, inhibited ecdysis, resulting in larval retention of the exuvial head capsule and the insect’s death. Similarly, larvae of the greenhouse whitefly exhibited 100% mortality when fed on *Ajuga reptans* plants. High levels of the three major phytoecdysteroids, 20-hydroxyecdysone (ecdysterone), makisterone A and cyasterone, have been found in several plants, including *Ajuga* (Tomás et al. 1992; Wessner et al. 1992; Dinan 2001; Coll and Tandrón 2005; Castro et al. 2008, 2011; Grace et al. 2008; Sun et al. 2012; Lva et al. 2014; Guibout et al. 2015; Taha-Salaime et al. 2019), quinoa and spinach (Dinan 1995b, 2001). An extract of 20-hydroxyecdysone and cyasterone from *A. iva* showed high activity against *Oligonychus perseae* (Kubo and Klocke 1983; Aly et al. 2011); a dose of 5 μg/ml of pure extracted *A. iva* ecdysterone significantly reduced fecundity, fertility and survival of this pest, while commercial 20-hydroxyecdysone at the same dose had lesser effects (Aly et al. 2011).

In addition to phytoecdysteroids, species of the genus *Ajuga* also contain the bioactive compounds clerodane diterpenes (include clerodanes) and iridoid glycosides (Camps and Coll 1993). Clerodanes (diterpenoids) are a large group of C20 terpene compounds derived from geranylgeranyl diphosphate and biosynthesized through the deoxyxylulose phosphate pathway in the cytoplasm, mostly in the leaves and stems of the Lamiaceae and Asteraceae families (Hussain et al. 2012). Clerodin was originally isolated from *Clerodendrum infortunatum* L. (Lamiaceae), and has potential as a natural pesticide due to its insect antifeedant and repellent activities (Pereira and Gurudutt 1990; Kubo et al. 1991; Coll and Tandrón 2005; Koul 2016; Li et al. 2016). Koul (2016) showed that the most active compounds, dihydroclerodin and clerodin hemiacetal, from *Caryopteris divaricata* exhibit 100% antifeedant activity at 50 ppm. These clerodanes were deadly to *Spodoptera litura* larvae.

We previously identified and quantified high contents of three phytoecdysteroids and two clerodanes in *A. iva* growing in Israel (Taha-Salaime et al. 2019). We hypothesized that crude extract of *A. iva* leaves that includes the three phytoecdysteroids (20-hydroxyecdysone, makisterone A and cyasterone), which specifically interfere by controlling molting, and are responsible for the metamorphosis and antifeedant activities in insects, might be a promising pest-control agent. We evaluated the efficiency of *A. iva* extracts (containing phytoecdysteroids and clerodanes) at reducing the damage caused by *S. littoralis* larvae by addressing the following questions: Does *A. iva* crude leaf extract affect *S. littoralis* larvae? Do phytoecdysteroids isolated from the crude leaf extract and commercial standards have different effects on the larvae? Do phytoecdysteroids have a direct effect on the larvae’s gut?

## Materials and Methods

### Plants and insects

*A. iva* plants were collected in April 2014 from a wild population in the Negev, southern Israel, and then cultivated and acclimated in an open field at Newe Ya’ar Research Center. Young and mature leaves and stems of fresh plants were collected after blooming (July– November) and oven-dried at 55°C for 3–4 days, then homogenized to a fine powder prior to extraction. The first and third instars of *S. littoralis* larvae used for the bioassays were from Murad Ghanim’s laboratory, Department of Entomology, Agricultural Research Organization (African cotton leafworm colony) reared on castor bean leaves.

### Extraction and purification of phytoecdysteroids

*A. iva* crude extracts were prepared according to our recently published procedure (Taha-Salaime et al. 2019). Leaf and stem powder were pooled (24 g) and soaked in 240 ml of 100% MeOH, sealed and homogenized with shaking (2500 rpm) for 1 h. The extract was then centrifuged (112 *g)* for 10 min, filtered and concentrated under vacuum. The final filtered methanol solution was analyzed by liquid chromatography-time of flight-mass spectrometry (LC-TOF-MS) and dried in a chemical vaporizer for 5 days. For purification of phytoecdysteroids from the crude extract, leaf and stem powder (100 g) was soaked in 300 ml methanol and homogenized. The filtered extract was vacuum-concentrated and treated with H_2_O to give 30% aqueous methanol. This solution was extracted as previously described (Aly et al. 2011).

### Identification phytoecdysteroids and clerodanes

LC-TOF-MS analysis was used to identify and confirm the presence of phytoecdysteroids and clerodanes in three concentrations of *A. iva* crude leaf extract (50, 100 and 250 µg/µl). We analyzed the profile of phytoecdysteroids and clerodanes in the *A. iva* crude leaf extract before each test for biological activity. Extracts of the plant material (1 µl) were injected into an Agilent 1290 Infinity Series liquid chromatograph coupled with an Agilent 1290 Infinity DAD and Agilent 6224 Accurate Mass TOF mass spectrometer (Agilent Technologies, Santa Clara, CA, USA) (Dinan 1989). Thin-layer chromatography (TLC) was used to separate the components into well-defined spots. The crude leaf extract, the pure isolated compounds (20-hydroxyecdysone [ecdysterone], makisterone A and cyasterone) and a commercial ecdysterone sample were applied to silica gel GF-254 plates (0.25 mm; 20 × 20 cm) as described in Aly et al. (2011).

### Biological activity of *A. iva* crude leaf extract against *S. littoralis*

To assess effects on the larvae, mature castor bean leaves were smeared, using a paint brush, with aqueous *A. iva* crude leaf extract (24 g of dried pooled leaves and stems dissolved in 240 ml MeOH, 1:10) and Tween 20 (1.5 mg). The leaves were dried in a chemical hood for 2 h. Then 10 first instar *S. littoralis* larvae were placed on 1 treated castor bean leaf in a petri dish and allowed to feed for 3 days in a climate-controlled room at 25°C. Control leaves were similarly smeared with double-distilled water (ddH_2_O) and Tween 20. The larvae were exposed to three concentrations of crude leaf extract (50, 100 and 250 µg/µl), one concentration per treatment. After preparing the *A. iva* crude leaf extract, the methanolic extract was dried in a chemical hood; 1.2 g dried extract powder was dissolved in 4.8 ml ddH_2_O and Tween 20 (0.5 mg/ml) for the 250 µg/µl concentration; 0.83 ml of the high concentration (250 µg/µl) extract was dissolved in 3 ml ddH_2_O to obtain the 100 µg/µl concentration; and 0.67 ml of the 100 µg/µl solution was dissolved in 3 ml ddH_2_O to obtain the 50 µg/µl concentration. At the end of the experiment, larval mortality was compared to that of controls. Data in this experiment represent the results of 11 replicates (10 larvae/replicate). Differences are reported as percent mortality of first instar larvae after feeding on the three concentrations of *A. iva* crude leaf extract using a t-test and significance was determined by t-test.

For the third instars, 1 or 10 larvae were fed on 1 treated caster bean leaf for 4 or 8 days in a climate-controlled room at 25^°^C (more freshly treated leaves were provided after 4 days of feeding to avoid feeding on decayed leaves). Four different treatments were tested, where larvae were fed on castor bean leaves treated with: (1) 250 µg/µl *A. iva* crude leaf extract for 4 days, with a freshly treated leaf for 4 more days; (2) the same treatment as (1) with 250 µg/µl of a fractionated mixture of three phytoecdysteroids from *A. iva* leaf extract; (3) *A. iva* crude leaf extract for 4 days, and then a control castor bean leaf treated with ddH_2_O for the next 4 days; (4) a control castor bean leaf for 4 days and then *A. iva* crude leaf extract for the next 4 days. In parallel, control leaves were smeared with ddH_2_O and Tween 20 for 8 days. We recorded the different reactions of *S. littoralis* larvae in all treatments after 4 days, and if some larvae can recover again if provided control castor bean leaf larvae (first were fed on treated leaf), or were adversely affected and dead when provided a treated castor bean leaf after 4 days (first were fed on control leaves. In another experiment, third instar larvae were exposed to three concentrations of *A. iva* crude leaf extract (50, 100 and 250 µg/µl) for 4 days. At the end of the experiment, we recorded larval survival and relative growth rate (RGR) as (ln *W*_2_ – ln *W*_1_)/ (*t*_2_ – *t*_1_), where *W*_1_ and *W*_2_ are weights at times *t*_1_ and *t*_2_ (n = 20 replicates). In each treatment, pupation rate was evaluated for an additional 15 days. RGRs were compared using a mixed-model ANOVA (repeated measures ANOVA until day 4 and two-way ANOVA from day 4 to the end of the experiment). Survival rates were analyzed using a Friedman test, since the data for larval survival did not follow a normal distribution. Data for larval weight gain (LWG) are the result of 10 replicates (10 larvae in each replicate). Comparisons of LWG were assessed using a repeated measures ANOVA. Statistical significance was reported at *p* < 0.05. Error bars in all graphs represent the standard error of the mean (SEM), and significance is indicated in each experiment. All statistical analyses were performed with IBM SPSS software v.20 for Windows (IBM, Armonk, NY, USA).

### The effect of *A. iva* crude leaf extract and purified phytoecdysteroid mixture on larval gut of *S. littoralis*

To examine the gut morphology of *S. littoralis* larvae, we used DAPI and phalloidin staining. Larvae fed on castor bean leaves treated with *A. iva* crude leaf extract and controls (leaves treated with water) were tested after 7 days of treatment. Guts were dissected in phosphate buffered saline (1X PBS), then fixed in 4% paraformaldehyde in 1X PBS for 30 min, washed in 0.1% Triton X-100 for 30 min, washed three times in PBS Tween-20 (PBST) (https://www.usbio.net/protocols/phosphate-buffered-saline-tween-20), incubated in 0.1% phalloidin in PBST for 30 min, washed three times with PBST and mounted whole in 0.1% DAPI in hybridization buffer (20 mM Tris-HCl, pH 8.0, 0.9 M NaCl, 0.01% w/v sodium dodecyl sulfate, 30% v/v formamide). Changes in actin fibers and nuclei were visualized under a Leica confocal microscope (Leica SP8 and Olympus IX 81 confocal laser scanning microscope).

## Results

### Identification of natural phytoecdysteroids from *A. iva* using TLC analysis

*A. iva* crude leaf extract was subjected to flash chromatography on a silica gel (TLC), yielding three individual isolated compounds (20-hydroxyecdysone, makisterone A and cyasterone). The retention factor (Rf) values, i.e., the distance migrated over the total distance covered by the solvent, of the phytoecdysteroid spots were similar to those of the respective commercial ecdysteroids (Fig. 1).

**Fig. 1.**
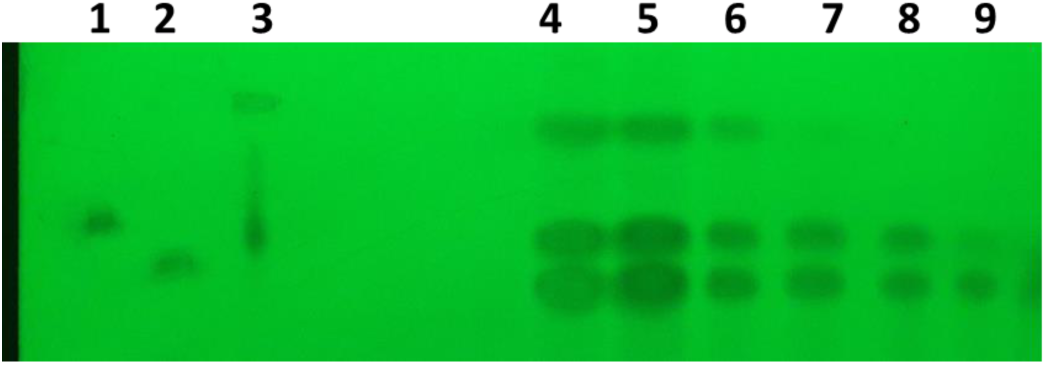
Identification of *A. iva* phytoecdysteroids by TLC. TLC plate shows the separation of three phytoecdysteroids (20-hydroxyecdysone, makisterone A and cyasterone) (4–6), commercial ecdysterone standards (1–3): makisterone (1), 20-hydroxyecdysone (2) and cyasterone (3). Fractions (7–9) show the presence of only 20-hydroxyecdysone and cyasterone. Fractions 4–6 were used in the bioassays

### The effect of *A. iva* crude leaf extract on first instar larval survival

First instar *S. littoralis* larvae showed a significant increase in mortality (25, 65, 85%) after feeding on the three concentrations of *A. iva* crude leaf extract (50, 100 and 250 µg/µl, respectively), compared to the control (treated with water, 5%) (Fig. 2a).

**Fig. 2.**
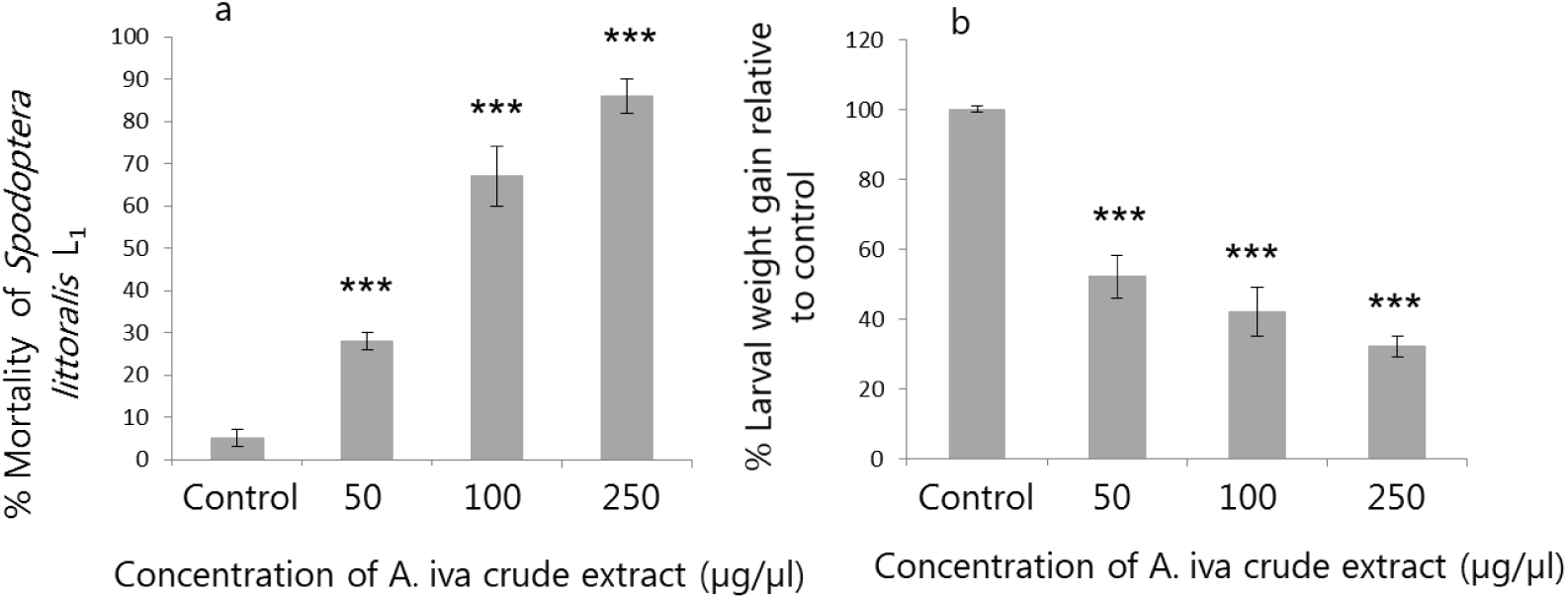
Effect of different concentrations of *A. iva* crude leaf extract on *S. littoralis* first instar (L_1_) larval (n = 110) mortality (mean ± SEM) (a), and larval weight gain (%) (mean ± SEM) of *S. littoralis* third instar (L_3_) larvae (b). Asterisks above columns indicate significant difference (*p* ≤ 0.05) by t-test (t_108_ = 6.105, 4.308 and 3.220 for 50, 100 and 250 µg/µl, respectively); *p* < 0.001 for all treatments, Levene’s test *p* = 0.326 (a), and by repeated measures ANOVA (F_3.104_ =20.334, 17.246 and 13.007 for 50, 100 and 250 µg/µl, respectively); *p* < 0.001, Mauchly’s test *p* = 0.152 (b) between treatments and the control

### The effect of *A. iva* crude leaf extract on third instar larval survival and development

Third instar *S. littoralis* larvae fed on crude leaf extract (50, 100 and 250 µg/µl) showed reduced LWG (F_3.104_ = 20.334, 17.246 and 13.007, respectively, *p* < 0.001; Fig. 2b) compared to the control.

All concentrations of crude leaf extract significantly decreased (*p* < 0.001) larval RGR compared to the normally developing larvae on the control diet (Fig. 3a, arrow). Reduced RGR was recorded as early as 2 days into the experiment. With the highest concentration of crude leaf extract, RGR decreased by 0.05 and 0.20 g/g day on days 6 and 8, respectively, compared to the control (F_3.16, 18_ =12.641, *p* < 0.001; Fig. 3a). All concentrations of *A. iva* crude leaf extract significantly reduced third instar larval survival after 11 days (*X*^*2*^_3_ = 6.221, *p* = 0.038; Fig. 3b). Whereas all larvae survived on the control leaves, the effect of the crude extract was apparent after 3 days. In fact, none of the treated larvae survived more than 8 days for the highest concentration of crude leaf extract and 10 days for the other concentrations (Fig. 3b).

**Fig. 3.**
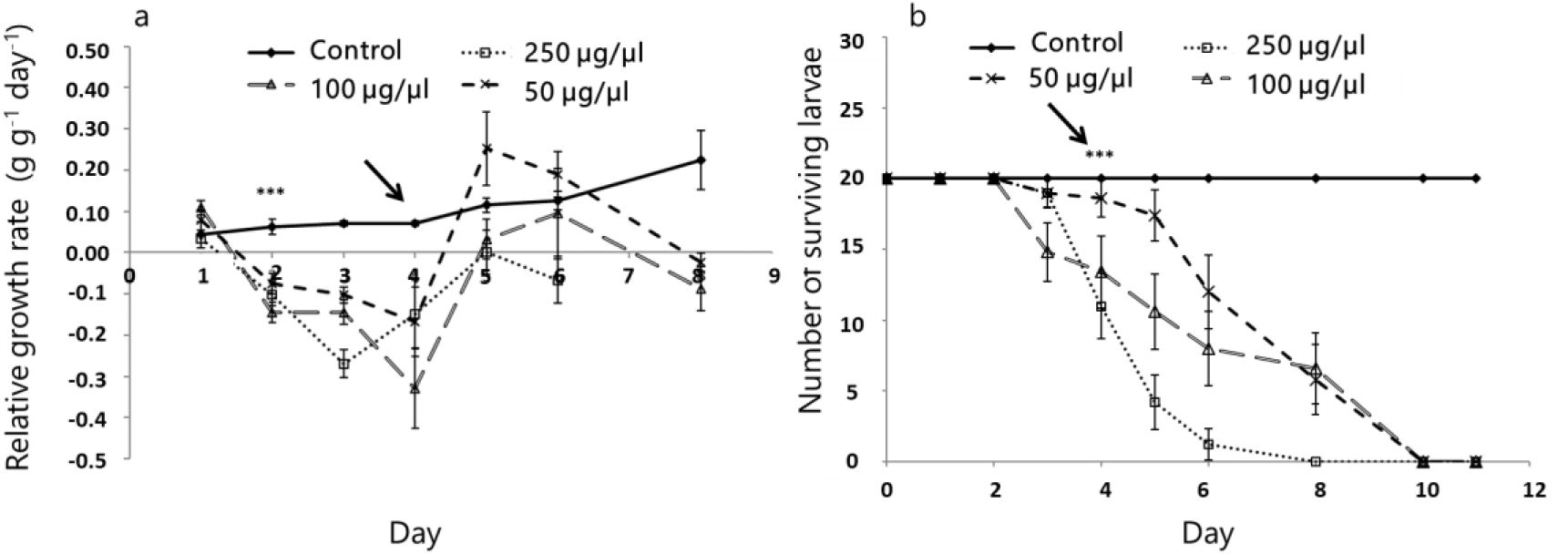
Effect of different concentrations of *A. iva* crude leaf extract on *S. littoralis* third instar larval relative growth rate (mean ± SEM) (a) and survival (b); n = 20. Asterisks above points indicate significant difference (*p* ≤ 0.05) between treatments and the control

In addition, when *S. littoralis* larvae were first fed on control leaves for 4 days and then on leaves treated with *A. iva* crude leaf extract for an additional 4 days, their RGR was affected by feeding on the crude leaf extract after day 5 of the experiment (F_3.16, 18_ =7.310, *p* < 0.001; Fig. 4a), and continued to decrease until the end of the experiment, with no surviving larvae (*X*^*2*^_3_ = 9.282, *p* = 0.021; Fig. 4b).

**Fig. 4.**
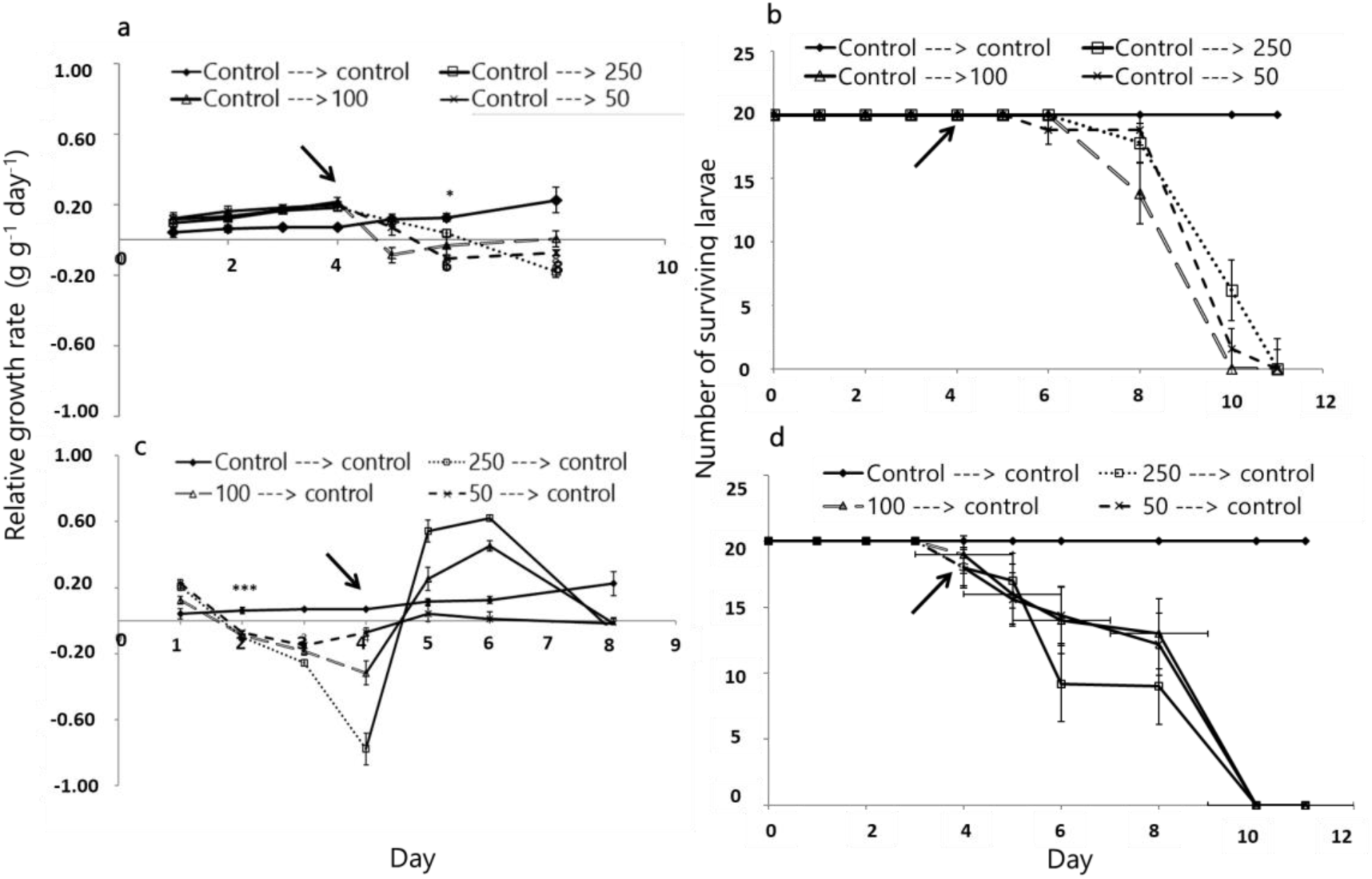
Effect of *A. iva* crude leaf extract on *S. littoralis* third instar larval relative growth rate (mean ± SEM) and survival. Larval relative growth rate (a) and survival (b) when fed on control leaf (treated with water) until day 4, and then fed on leaves treated with crude leaf extract at three concentrations until the end of the experiment (a). Relative growth rate (c) and survival (d) of the larvae after feeding on crude leaf extract at three concentrations until day 4 and then control leaves until the end of the experiment; n = 20. Asterisks above points indicate significant difference (*p* ≤ 0.05) between treatments and the control. Arrow points to day 4

The same result was obtained when the order of the treatments was reversed (Fig. 4c, d). When the larvae were first fed on leaves treated with *A. iva* crude extract for 4 days and then fed on control leaves (for an additional 4 days), a significant decrease in RGR was obtained on days 2–4 (F_3.16, 18_ = 4.595, 3.608 and 8.113, *p* = 0.034, 0.02 and 0.001 for 50, 100 and 250 µg/µl crude leaf extract, respectively; Fig. 4c). Moreover, castor bean leaves treated with 250 µg/µl of the mixture of the three fractionated and purified phytoecdysteroids from the *A. iva* crude leaf extract significantly reduced RGR (F_2.20, 18_ = 6.172, *p* = 0.001) compared to the control (Fig. 5a). Few larvae survived on the leaves treated with purified phytoecdysteroid fraction (*X*^*2*^_3_ = 11.305, *p* = 0.04) (Fig. 5b).

**Fig. 5.**
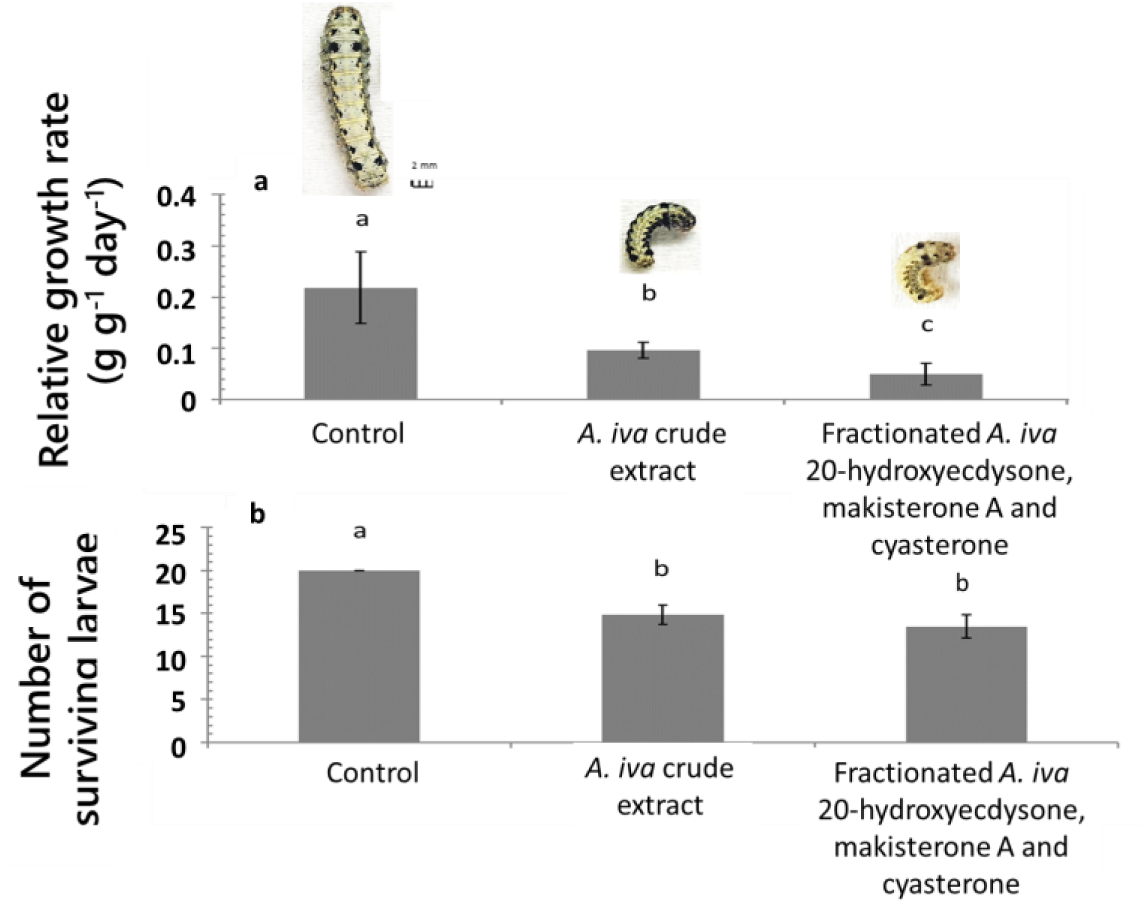
Effect of *A. iva* crude leaf extract (250 µg/µl), and of the three fractionated and purified phytoecdysteroids (250 µg/µl) on *S. littoralis* third instar larval relative growth rate (mean ± SEM) (a) and survival (b). The phytoecdysteroid fraction contained 20-hydroxyecdysone, makisterone A and cyasterone; n = 20. Development of *S. littoralis* larvae shown above the columns after 4 days feeding on control leaves, or leaves treated with 250 µg/µl *A. iva* crude leaf extract or 250 µg/µl of the three fractionated phytoecdysteroids. Different letters above columns indicate significant difference (*p* ≤ 0.05) between treatments and the control

Overall, larvae fed on castor bean leaves treated with 250 µg/µl *A. iva* crude leaf extract or 250 µg/µl of the phytoecdysteroid mixture lost weight, stopped growing and ultimately died (Fig. 5a, larvae depicted above columns).

### The effect of *A. iva* crude leaf extract and purified phytoecdysteroid mixture on larval gut of *S. littoralis*

Larval guts were stained with phalloidin, an actin-specific marker that binds to the interface between adjacent actin monomers in the F-actin polymer, and with DAPI, which stains the nuclei. Larvae feeding on 250 µg/µl *A. iva* crude leaf extract for 8 days had smaller nuclei with an abnormal shape—the nuclei moved to the edges of the cell and were thinner than normal (Fig. 6d–f). Phalloidin staining showed normal actin-filament organization in the control treatment (Fig. 6a–c). In contrast, in guts dissected from larvae treated with 250 µg/µl crude leaf extract or 250 µg/µl of the three phytoecdysteroids (20-hydroxyecdysone, makisterone A and cyasterone) isolated from the leaf extract, the actin filaments were smaller and their amount reduced (Fig. 6d–i). The damage observed in these experiments continued until the insects died. Overall, larvae exposed to crude leaf extract or its phytoecdysteroid fraction had less actin fibers and smaller, abnormally shaped nuclei.

**Fig. 6.**
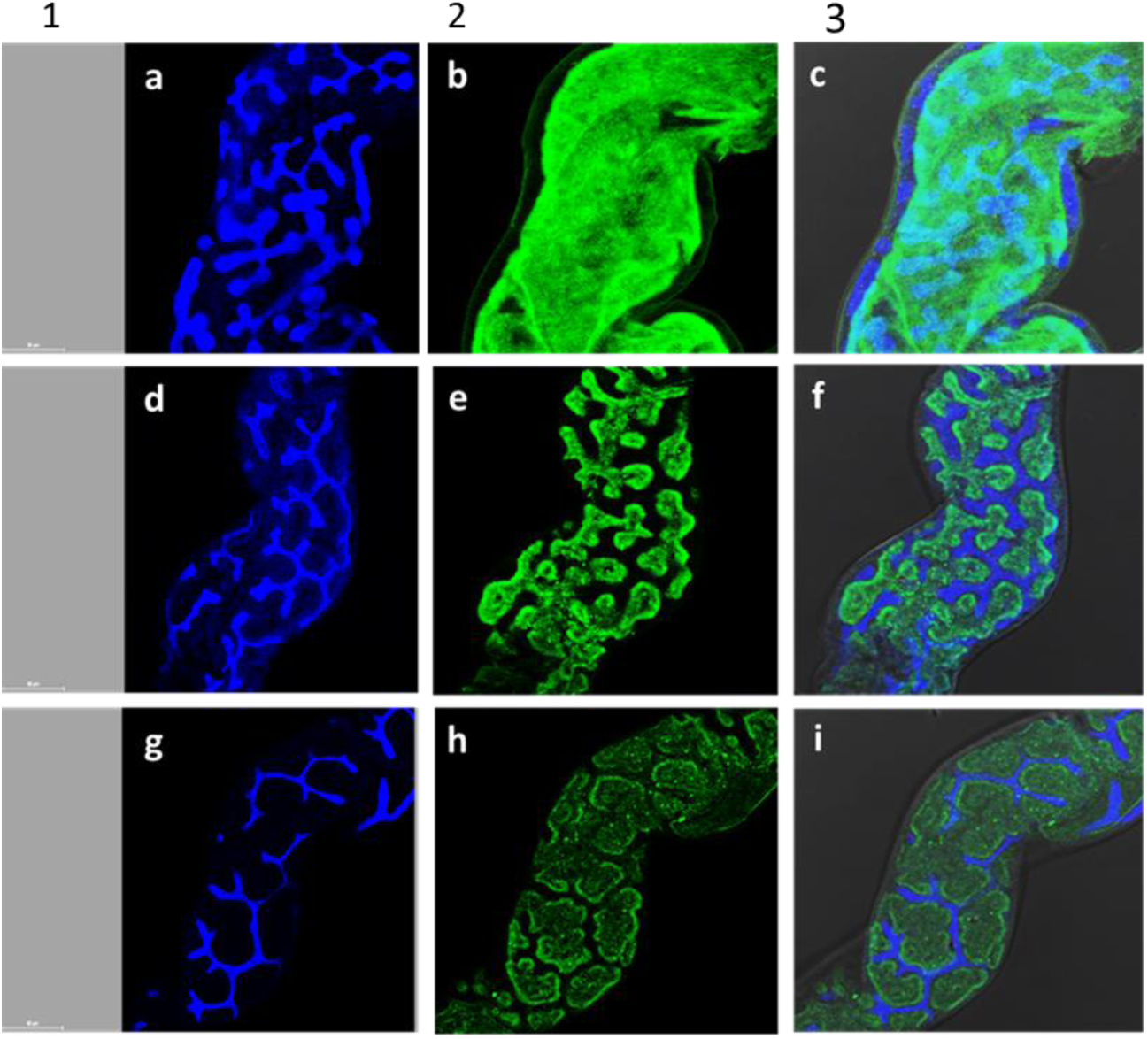
Gut morphology of *S. littoralis* third instar larvae after feeding on treated or non-treated leaves: control castor bean leaves treated with water (a–c), leaves treated with *A. iva* crude leaf extract (250 µg/µl) (d–f), and leaves treated with 250 µg/µl of three fractionated and purified phytoecdysteroids from *A. iva* leaf extract (20-hydroxyecdysone, makisterone A and cyasterone) (g–i) for 8 days (60 µm, respectively). Blue: DAPI staining of the nuclei under dark field (1); green: phalloidin staining of actin filaments under dark field (2), and double DAPI staining of the nuclei and phalloidin staining of actin filaments under dark field (3)

### Pupation of *S. littoralis* larvae following biological activity treatments

Larvae fed 250 µg/µl of crude leaf extract or 250 µg/µl of its phytoecdysteroid fraction for 8 days were unable to complete their development and pupate after 15 days (Fig. 7). Figure 7b and c shows the incomplete pupae obtained; the dying larvae had short limbs, small heads, decreased weight and only stomach and chest pupated, considered non-pupation. None of them completed their development to adult moths.

**Fig. 7.**
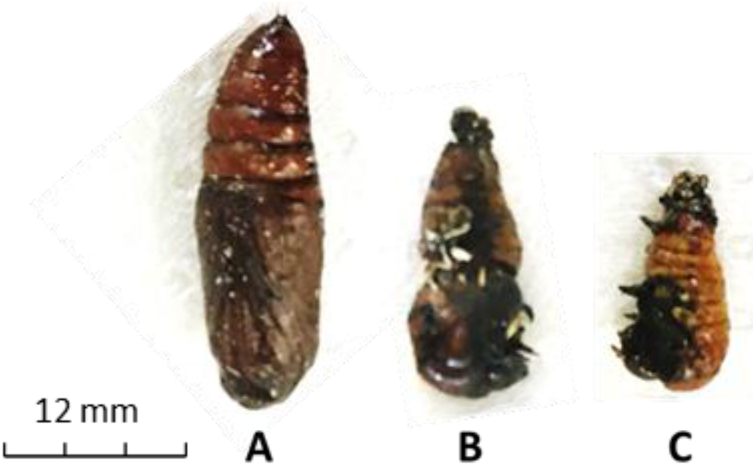
Metamorphosis of control and treated *S. littoralis* larvae. Pupation of *S. littoralis* larvae after 15 days of exposure (feeding for 4 days) on *A. iva* crude leaf extract. Control (treated with water) (a), 250 µg/µl *A. iva* crude leaf extract (b) and 250 µg/µl of three fractionated and purified phytoecdysteroids from *A. iva* leaf extract fractions (20-hydroxyecdysone, makisterone A and cyasterone) (c). Deficient development of pupation in (b) and (c) is due to lower levels of the ecdysteroids responsible for molting

## Discussion and Conclusions

As predicted by Taha-Salaim et al. (2019), we found that Israeli *A. iva* crude leaf extract and its fractionated phytoecdysteroids (20-hydroxyecdysone, makisterone A and cyasterone) significantly reduce the development and survival of *S. littoralis* larvae. These effects were pronounced throughout all larval developmental stages, including pupation. It has been shown that phytoecdysteroids negatively affect lepidopteran pests (Schmelz et al. 2002), whereas other insect species tolerate them (Schmelz et al. 2002; Taha-Salaime et al. 2019). We found that *S. littoralis* first and third instar larvae fed on *A. iva* crude leaf extract (50, 100 and 250 µg/µl) for 3 and 11 days, respectively, had increased mortality, reduced LWG and decreased RGR compared to the control treatment. Similarly, phytoecdysteroids from *A. iva* have been found to reduce the fertility and fecundity of *Bemisia tabaci* and *Oligonychus perseae* (Aly et al. 2011).

Our results are in agreement with previous studies suggesting that insect herbivores cannot develop and survive when fed on phytoecdysteroid-treated leaves. Ecdysteroids inhibited feeding of *Pieris brassicae* and *Mamestra brassicae* larvae when given at 200 mg/kg fresh weight in sucrose solution (Ma 1972), and inhibited drinking in *Dysdercus koenigii, Dysdercus fulvoniger* and *Spilostethus pandurus* adults at a concentration of 100 mg/kg (Schoonhoven and Derksen-Koppers 1973). Jones and Firn (1978) reported that ecdysone and 20-hydroxyecdysone deter feeding in *Pieris brassica* when incorporated above 5 mg/kg diet. Exogenous application of ecdysteroids was shown to be lethal to *Plodia interpunctella* and *Bombyx mori* larvae; ingestion of these compounds was toxic to the midgut epithelial cells (Tanaka and Yukuhiro 1999; Rharrabe et al. 2009; Wadsworth et al. 2014). In our study, *A. iva* crude leaf extract was most effective at the highest concentration applied, indicating a dose-dependent effect, in agreement with other studies (Tanaka and Takeda 1993). In contrast, *S. littoralis* was not deterred from feeding by 20-hydroxyecdysone at 50–70 mg/kg (Jones and Firn 1978).

In our study, LWG and RGR of *S. littoralis* larvae were affected by feeding on 250 µg/µl methanolic crude leaf extract dissolved in ddH_2_O, regardless of larval age, in agreement with recent research using a methanolic extract of *Ajuga remota* leaves containing cyasterone and ecdysterone, which disrupted the molting cycle in *Bombyx mori* and *Spodoptera frugiperda* (Kubo et al. 1981). Moreover, Slama et al. (1993) found that cyasterone and turkesterone are the most effective lepidopteran- and coleopteran-specific ecdysteroids, and ingesting the phytoecdysteroid 20-hydroxyecdysone caused death before and during *Bombyx mori* molting (Chou and Lu 1980).

In the current study, we did not fractionate clerodanes from *A. iva* crude leaf extract due to the difficulty involved in calibrating the protocol for fractionation, and to the high cost of commercial clerodane standards. Kubo (1993) conducted an artificial diet-feeding assay with the wheat aphid *Schizaphis graminum*, and showed that ajugasterone C (a clerodane) was 10-fold more potent as a feeding deterrent than 20-hydroxyecdysone, and 30-fold more potent than cyasterone (Kubo 1993).

In our study, we only used a few commercial standards because they are very expensive and are not feasible as a control treatment. We could not use them at the same concentrations as the applied treatment. Therefore, we conducted an experiment with a mixture of three commercial standards (ecdysterone, makisterone A and cyasterone) at a maximum concentration of 100 ppm each, which is very low compared to the concentration in the (crude leaf extract and phytoecdysteroid fraction treatments (250,000 ppm). *S. littoralis* larvae were fed on castor bean leaf treated with 100 ppm of the mixture for 8 days. No significant effect of the mixture on *S. littoralis* larvae was seen. Tanaka (1995) reported altered epidermal sensitivity to 20-hydroxyecdysone at 300 ppm ecdysone, higher than the standard concentration used in our study.

Based on the results, phytoecdysteroids may affect insects by interfering with their developmental stages, especially during metamorphosis (Chou and Lu 1980). In the present study, we observed suppressed pupation of *S. littoralis* due to the reduction in LWG; the larvae did not reach the threshold weight for pupation and they died because they could not complete their life cycle. The histological observations of the gut showed that *S. littoralis* is very sensitive to *A. iva* crude leaf extract and the mixture of the three fractionated phytoecdysteroids (20-hydroxyecdysone, makisterone A and cyasterone). The larval gut cells showed histolysis with clear signs of apoptosis. The gut epithelium showed massive deterioration, there was destruction of the microvilli of the columnar cells, and formation of vacuoles. In smaller larvae, mortality occurred during molting between instars, whereas in bigger larvae, most mortality was at the prepupal stage. Our results of gut cell destruction support the notion that the effect on pupation could be a consequence of disruptions in hormonal balance effected by internal levels of ecdysone. External ecdysteroid detoxification is one of the main ways in which insects overcome the toxic effects of these compounds. The transition from one stage in ovarian development to another, such as from previtellogenesis to vitellogenesis and then chorionogenesis, is governed by the actions of several pathways that respond to different titers of 20-hydroxyecdysone (Swevers and Iatrou 2003).

Feeding on phytoecdysteroids such as ecdysterone, polypodine B and ponasterone A induces ecdysial failure associated with the appearance of larvae having two head capsules and developmental anomalies during metamorphosis in *Acrolepiopsis assectella* (Arnault and Slama 1986). Since the reduced growth rate (Figs. 3, 4) suggests that larvae are adversely affected by ingestion of the crude leaf extract and of the mixture of three phytoecdysteroids from *A. iva*, we assume that the effect observed on LWG and RGR reflects another possible mode of action of phytoecdysteroids. Abnormal gut development can lead to reduced LWG, leading to mortality. In the present study, disruptions in *S. littoralis* gut morphology and disappearance of microfilament structures in actin (Fig. 6) could be a consequence of the phytoecdysteroid titers in *A. iva* crude leaf extract. Actin microfilaments in particular have been associated with the rounding and loss of adhesion that frequently occur with viral infection or transformation in response to secondary metabolites (Carley et al. 1981; Meyer et al. 1981), with the intracellular transport of viral structural proteins and viral particles (Bohn et al. 1986), with the budding process of many enveloped viruses (Mortara and Koch 1989), and with the assembly of virions in the cytoplasm (Jackson and Bellett 1989) and in the nucleus (Wang and Goldberg 1976).

In conclusion, our data suggest that the phytoecdysteroids and clerodanes of *A. iva* may be useful for the management of economically important insect pests such as *S. littoralis*, while reducing the risks to human health and the environment.

## Acknowledgments

This work was supported by the Israeli Ministry of Science and Technology (research grant no. 3-14496). We thank Dr. Rachel Davidovich-Rikanati and Alona Sheachter for their fruitful advice and comments.

